# Harmonic memory signals in the human cerebral cortex induced by sematic relatedness of words

**DOI:** 10.1101/2022.09.29.510214

**Authors:** Yasuki Noguchi

## Abstract

When we memorize multiple words simultaneously, semantic relatedness among those words assists memory. For example, the information of “apple”, “banana” and “orange” will be connected via a common concept of “fruits” and become easy to retain and recall. Neural mechanisms underlying this semantic integration in verbal working memory remain unclear. Here I used electroencephalography (EEG) and investigated neural signals when healthy human participants memorized five nouns semantically related (Sem trial) or not (NonSem trial). The regularity of oscillatory signals (8 – 30 Hz) during the retention period was found to be lower in NonSem than Sem trials, indicating that memorizing words unrelated to each other induced a non-harmonic (irregular) waveform in the temporal cortex. These results suggest that (i) semantic features of a word are retained as a set of neural oscillations at specific frequencies and (ii) memorizing words sharing a common semantic feature produces harmonic brain responses through a resonance or integration (sharing) of the oscillatory signals.

## Introduction

Verbal working memory (vWM) plays a critical role in various human behaviors such as reading, conversation, and inference, although its neural underpinnings remain controversial (Wilsch and Obleser, 2016; Oberauer et al., 2018; Pavlov and Kotchoubey, 2020). A hallmark of vWM is that its load can greatly change depending on a relationship among memory items (Bein et al., 2015). For example, when multiple words in a memory list are semantically associated (“apple”, “banana” and “orange”, etc.), the encoding and retention of those words is facilitated because they are integrated into a coherent concept (fruits) in the brain. In behavioral experiments, this is evidenced by high accuracy of a recognition task to judge whether a probe word (e.g. “dog”, normally presented in the end of a trial) matched any of the memory words or not. In contrast to this adaptive aspect (Schacter et al., 2011), integrating memory items also has a negative effect, sometimes inducing an erroneous response in the recognition task (Atkins and Reuter-Lorenz, 2011; Johns et al., 2012; Gatti et al., 2021a). If the probe is a lure word that is semantically related to memory items but has never appeared in the list (e.g. “pear”), typical participants falsely remember having seen the lure (called the “false memory” or “semantic interference”).

Many studies have investigated brain activity underlying the semantic integration and interference in memory (Martin et al., 2018; Reber et al., 2019; Zhu et al., 2019; Wing et al., 2020). Approaches of functional magnetic resonance imaging (fMRI) and transcranial stimulation revealed critical brain regions such as the anterior temporal cortex (Chadwick et al., 2016; Diez et al., 2017), prefrontal cortex (Atkins and Reuter-Lorenz, 2011), and cerebellum (Gatti et al., 2021b). However, it remained unclear how the semantic information is bound together as neural (electrical) signals in the human brain. Based on a close relationship between WM and oscillatory brain activities (Weiss and Mueller, 2012; Hanslmayr and Staudigl, 2014; Roux and Uhlhaas, 2014; Miller et al., 2018; Gehrig et al., 2019; Noguchi and Kakigi, 2020), here I use electroencephalography and test a hypothesis that a semantic integration in WM is represented as a harmony of neural rhythms (**Fig. 1**). In this model, semantic features of each memory word are maintained as a set of neural oscillations at specific frequencies. Retention of words sharing a common feature induces a resonance or integration of those oscillatory signals, generating a dominant frequency (or frequencies) in a power spectrum (**Fig. 1A**). This would produce a regular (more harmonic) neural waveform with a limited set of frequencies, forming a strong memory representation resistant to degradation by neural noises (irregular waveforms).

**Figure 1.**
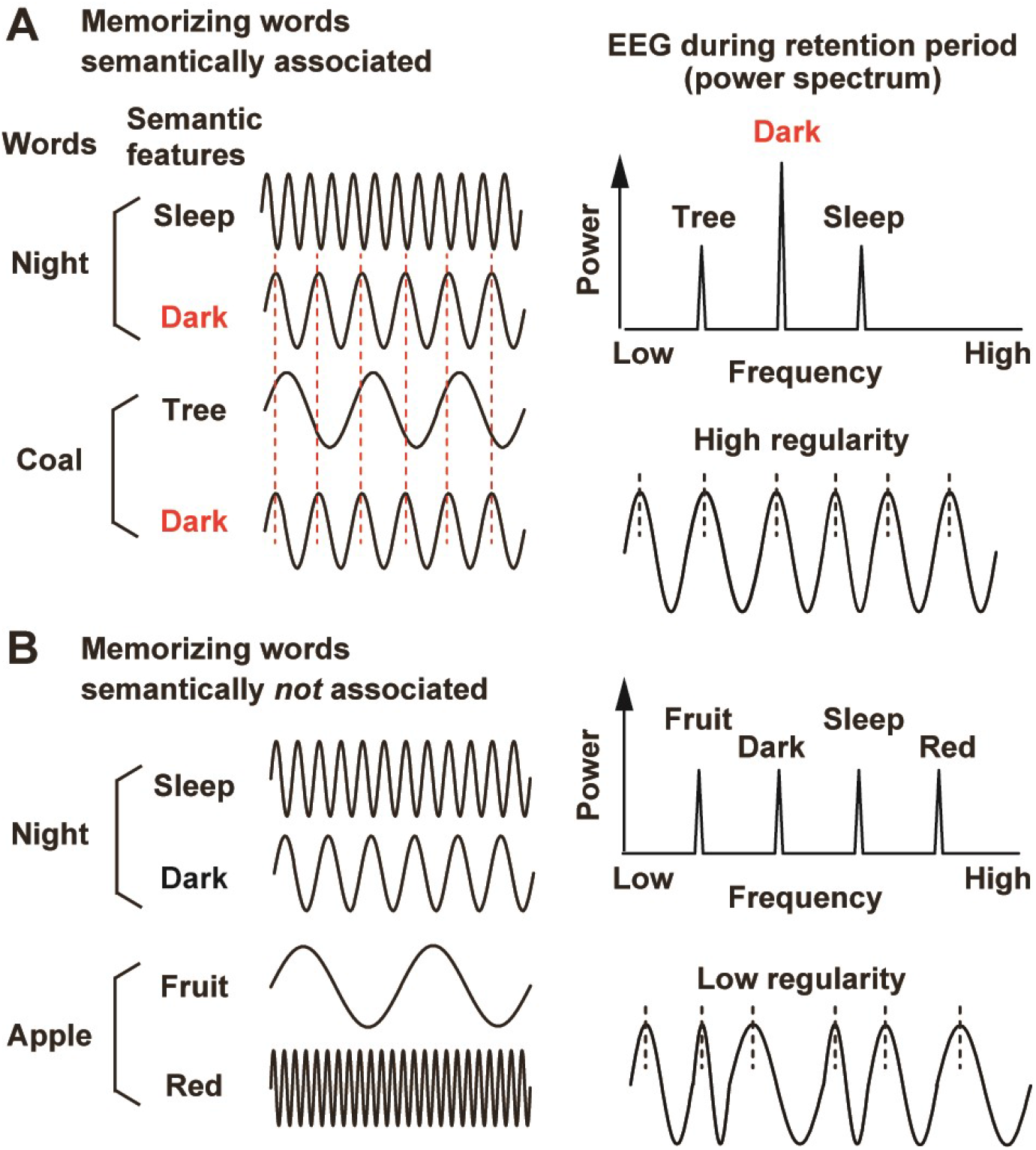
Scheme. For simplicity, here I assume that each semantic feature (e.g. sleep) of a word (e.g. night) is memorized as neural oscillation at a specific frequency. (**A**) Memorizing two words (e.g. night and coal) with a common feature (dark) induces a resonance or sharing of the oscillatory signal across the words, generating a regular EEG waveform characterized with a dominant frequency in a power spectrum (right panels). (**B**) Memorizing words without a common semantic feature induces no resonance or sharing of the oscillatory signal, producing an irregular EEG waveform composed of a number of different frequencies (like a white noise).

As shown in **Figure 2A**, participants in the present study attended to either left or right visual field and memorized five words sequentially presented. Neural activity during a retention period after the 5th word (delay 5 or D5) was compared between when the five words were semantically associated (Sem trial) or not (NonSem trial). If an across-word integration takes place as a neural harmony in the brain, this would be observed as higher regularity of EEG waveforms in Sem than NonSem trials.

**Figure 2.**
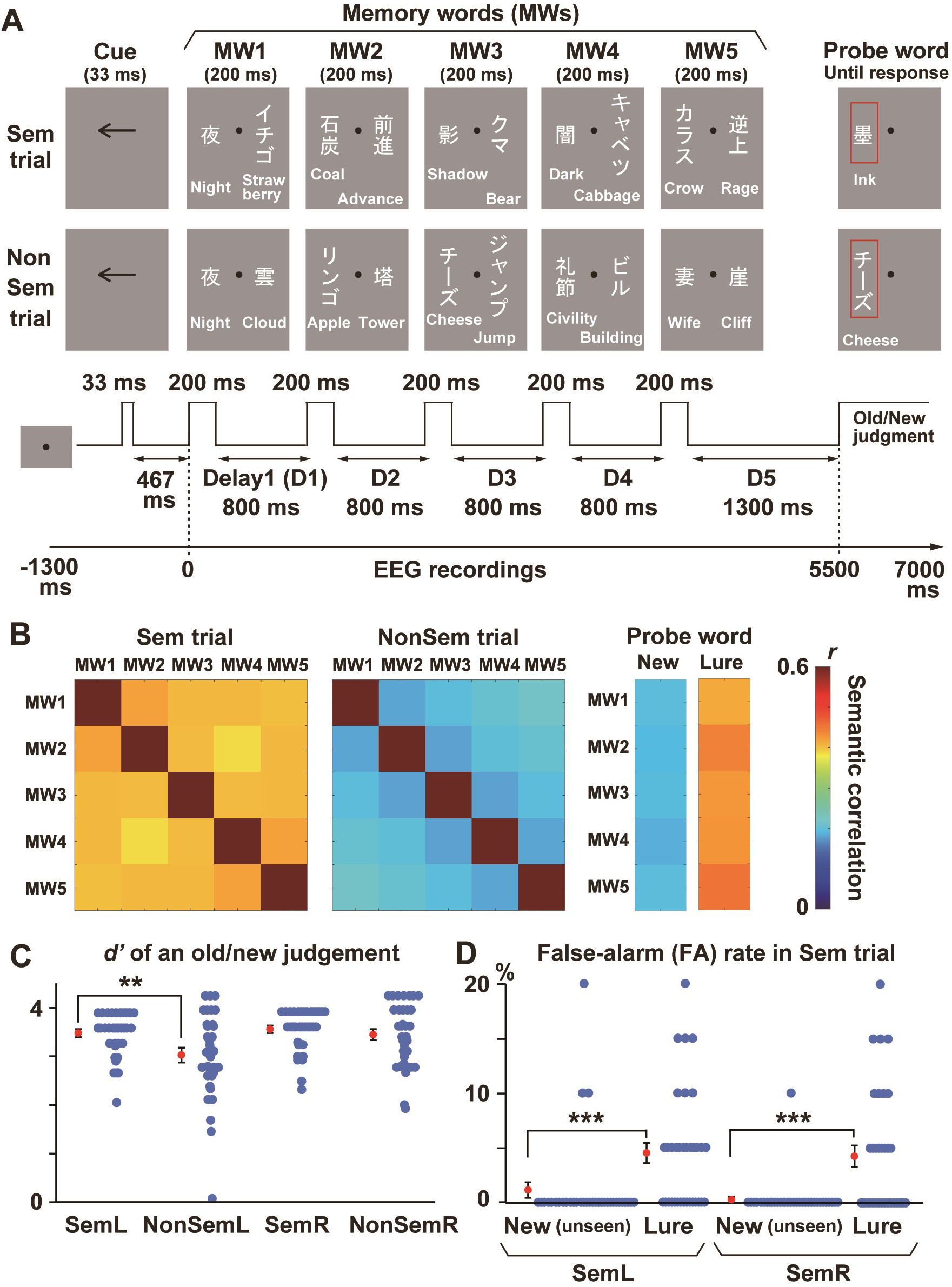
Experiment 1. (**A**) Each trial consisted of five screens (1 sec/screen, containing two words for each), followed by a probe word marked with a red rectangle. Participants memorized 5 words in a visual field indicated by an arrow cue (memory words or MWs). They pressed one button if the probe matched any of the five MWs (“old” response) and pressed another if not (“new” response). Neural activity during a retention period after the 5th word (D5) was compared between when the MWs were semantically associated (Sem trial, upper panels) or not (NonSem trial, lower panels). (**B**) Semantic relatedness among five MWs in Sem (left) and NonSem (middle) trials. The relatedness between two words was computed as a correlation between semantic vectors (1 × 300) of those words. Semantic correlations between MWs and two types of probe words in Sem trials (unrelated new words and lure words, see text) are also shown in the right panels. (**C**) The d-prime (*d’*) computed from hit and false-alarm (FA) rates in the old/new judgment task. Participants showed a better performance (higher *d’*) in Sem than NonSem trials only when they memorized MWs in the left visual field (Retain-Left conditions, SemL and NonSemL). (**D**) False memory. Higher FA rates to lure probes than unseen-new probes were observed both in SemL and SemR trials. Blue and red dots show individual data and an across-participant average, respectively. Error bars denote standard errors. ** *p* < 0.01, *** *p* < 0.001.

## Materials and Methods

### Participants

Thirty-four healthy subjects (native speakers of Japanese) participated in Experiment 1 (17 females, age range: 18 - 42). This sample size (34) was determined by a power analysis using G*Power 3 (Faul et al., 2007). The type I error rate and statistical power were set at 0.05 and 0.80, respectively. An effect size was assumed to be middle (0.5) (Cohen, 1988) because I found no previous study having the same goal as the present one. Data of one subject (female) were excluded from analysis due to excessive noise in EEG waveforms and thus replaced by data of an additional participant (female). Laterality quotients (LQs) measured by the Edinburgh Handedness Inventory (Oldfield, 1971) showed that all participants were right-handed (mean: 84.76, range: 11.11 - 100) but two (−11.11 and -17.65). Thirty healthy subjects participated in Experiment 2. Data of three participants were excluded from analyses because of a technical problem (loss of EEG data, N = 1) and excessive noise (N = 2), resulting in 27 participants in a final dataset (11 females, mean LQ: 80.28). All participants had normal or corrected-to-normal visual acuity. After the nature of the study had been explained, I received informed consent from each participant. All experiments were conducted in accordance with regulations and guidelines approved by the ethics committee of Kobe University, Hyogo, Japan.

### Task (Experiment 1)

All visual stimuli were generated with the Matlab Psychophysics Toolbox (Brainard, 1997; Pelli, 1997) and presented on a CRT monitor (refresh rate: 60 Hz). Each trial started with a black fixation point (0.18 × 0.18 deg) over a gray background for 800 ms. This was followed by a cue stimulus (an arrow pointing leftward or rightward, length: 1.34 deg, duration: 33 ms) over a central field to direct attention of participants (**Fig. 2A**). After another fixation period (467 ms), participants viewed two Japanese words (nouns), one in left and another in right visual fields, presented simultaneously for 200 ms (memory-word screen). Those words consisted of 1-5 white Japanese letters (Kana and Kanji characters) vertically arranged. A size of each letter was 1 (H) × 1 (V) deg, and a center-to-center distance between the fixation point and word was 1.25 deg. A total of five screens (ten words) were sequentially presented with an inter-screen interval (delay) of 800 ms. After the last (5th) delay of 1300 ms, the trial ended with a probe word (marked by a red rectangle, always shown in the cued visual field). Participants were asked to memorize five words in the cued hemifield (memory words or MWs) and performed an old/new judgment on the probe word. They pressed one button if the probe matched any of the five words they retained (“old” response) but pressed another if not (“new” response). They were also instructed to ignore five words presented in an uncued hemifield. No time limitation was imposed for this old/new judgement.

An effect of semantic integration on EEG waveforms was measured by a comparison of Sem and NonSem trials. In Sem trials, five MWs in the cued hemifield were semantically related (e.g. “piano”, “band”, “melody”, “concert”, and “jazz”), while they were not in NonSem condition (e.g. “curtain”, “jazz”, “rose”, “pencil”, and “kitchen”). Importantly, words in both conditions were taken from the same list of 300 words so that total visual inputs across all trials were balanced between Sem and NonSem. A combination of cued hemifield (left/right) and semantic relatedness (Sem/NonSem) produced four types of trials. Trials with a leftward cue and related MWs were called as SemL, while those with a rightward cue and unrelated MWs were called as NonSemR. An experimental session contained 60 trials in which those four types of trials (15 for each) were intermixed in a random order. A whole experiment consisted of four sessions.

Each of the four conditions (SemL, SemR, NonSemL, and NonSemR) had 60 trials in total. In SemL and SemR, one of five MWs was shown as a probe in 20 out of the 60 trials (old-probe trial, e.g. “jazz” in the above case). Serial positions (1 - 5) of the probe word were balanced (four trials for each position) to confirm the primacy and recency effects. A probe word not included in MWs was shown in another 20 trials (new-probe trial, e.g. “steak”). The new probe was either taken from five words in the uncued hemifield in the same trial (unattended new probe, 10 trials) or totally new (unseen new probe, 10 trials). In the other 20 trials, I showed a probe that was semantically related to the MWs but had never appeared (lure probe, e.g. “rhythm”). In NonSemL and NonSemR, 30 trials had an old probe and the other 30 trials had a new probe (unattended new probe in 20 trials and unseen new probe in 10 trials).

### Analysis of behavioral data

Behavioral data were analyzed using the signal detection theory. For each of the four conditions, I computed a hit rate in which participants answered “old” to an old probe and a false-alarm (FA) rate in which they answered “old” to a new probe. A measure of sensitivity (*d’*) was computed (Macmillan and Kaplan, 1985) using the equation

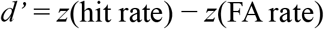

where *z* denotes the inverse cumulative normal function. If semantic relatedness across MWs facilitated the old/new judgment, this would be observed as higher *d’* in Sem than NonSem conditions.

Semantic integration, however, is also known to exert a negative influence when the probe is related to memory words (false memory). I examined this point by comparing the FA rates between the lures and new (unseen) probes of Sem condition. False memory would be indexed by a higher FA rate to lures than that to new probes.

### Stimuli (words)

MWs in each trial were taken from a list of 300 Japanese nouns prepared for the present study. Most words in this list were made by translating English words in the Deese–Roediger–McDermott (DRM) list (Roediger et al., 2001) into Japanese, although some words were arranged with procedures of Miyaji and Yama (Miyaji and Yama, 2002). A list of five MWs in Sem trial was determined based on the DRM list (e.g. “piano”, “band”, “melody”, “concert”, and “jazz”), while a list in NonSem trial was made by choosing five words unrelated to each other. Each word was used twice per condition to generate 240 trials in total. The validity of those lists was checked by computing semantic correlations among five MWs. I obtained a semantic vector (1 × 300) for each of the 300 words from fastText library (https://fasttext.cc/). Semantic relatedness between two MWs was measured as a correlation coefficient of those vectors. As shown in **Figure 2B**, semantic correlations among five MWs (averaged across all trials) were 0.39 – 0.42 in Sem and 0.18 – 0.22 in NonSem. I also confirmed that correlations between MWs and lure probes (0.41 – 0.45) were higher than those between MWs and new probes (0.19 – 0.20).

Other linguistic factors such as phonological variations were also controlled. I measured a within-trial variation of phonological factor by analyzing Japanese mora of each word (Chen et al., 2016). For each trial, a total number of moras used over five MWs was computed, with a mora shared by more than two MWs was counted as one (phonological variation). No significant difference was observed (*t*(238) = 0.42, *p* = 0.67, Cohen’s *d* = 0.05) between Sem (12.87 moras) and NonSem trials (12.98 moras).

### EEG measurements and analyses

Neural activity was recorded with the ActiveTwo system by Biosemi (Amsterdam, Netherlands). I measured EEG signals at 32 points over the scalp; FP1, FP2, AF3, AF4, F7, F3, Fz, F4, F8, FC5, FC1, FC2, FC6, T7, C3, Cz, C4, T8, CP5, CP1, CP2, CP6, P7, P3, Pz, P4, P8, PO3, PO4, O1, Oz, and O2 (**Fig. 3A**). Data were recorded with a sampling rate at 2,048 Hz and an analog low-pass filter of 417 Hz. Preprocessing of EEG data were performed with the Brainstorm toolbox (Tadel et al., 2011) for Matlab. I first removed a fixed-frequency noise from power line (60, 120, and 180 Hz) using a notch filter. A band-pass filter of 0.1 - 200 Hz was also applied to eliminate low- and high-frequency noises. For the data of six participants, a band-pass filter of 0.5 – 200 Hz was used because their data contained much low-frequency noise presumably caused by body movements. All data were then referenced with an average potential over the 32 electrodes. I segmented EEG waveforms into each trial (epoch range: -1300 to 7000 ms relative to an onset of the 1st MW screen) and classified them into the 4 conditions. Waveforms with a max-min amplitude larger than 150 μV at -700 to 5500 ms were excluded from analyses. Numbers of trials that remained after the rejection were 44.15 (SemL), 42.97 (SemR), 44.59 (NonSemL), and 43.85 (NonSemR). A two-way ANOVA of Sem/NonSem × L/R indicated no main effect or interaction (*F*(1,33) < 2.77, *p* > 0.10, *η*^2^ < 0.078 for all).

**Figure 3.**
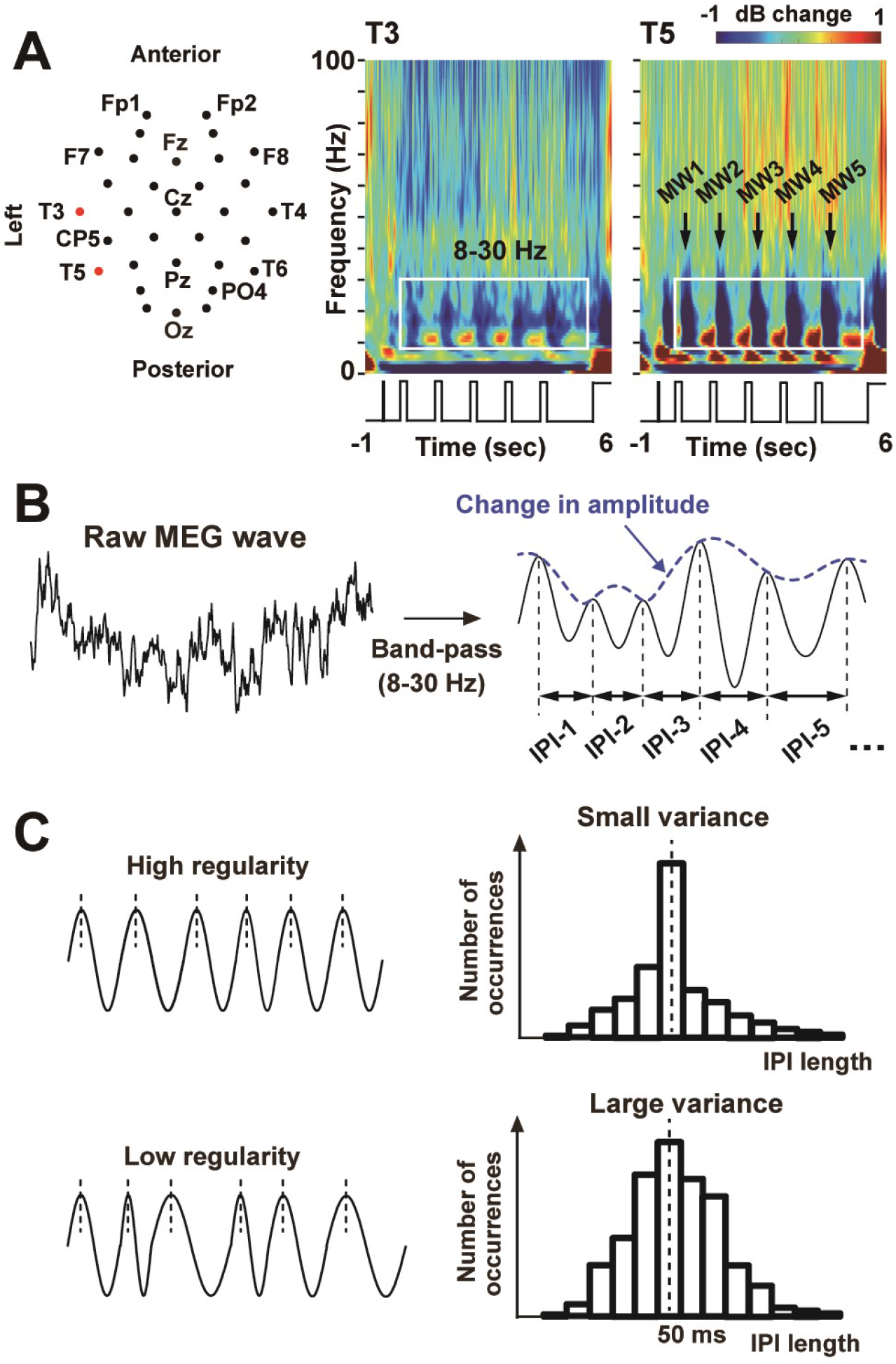
Measurements and analyses of electroencephalography (EEG) data. (**A**) Two-dimensional layout of 32 EEG sensors. Time-frequency power spectra (decibel power changes from a pre-cue period, -800 to -500 ms) at two sensors over the left temporal cortex (T3 and T5) are also shown. Prominent power changes throughout the five retention periods (D1 - D5) are seen in alpha-to-beta band (8-30 Hz) indicated by a white rectangle. (**B**) Three measures of oscillatory signals. I first extracted waveforms at the alpha-to-beta band with a band-pass filter. Changes in amplitude was measured as an envelope of the filtered waveform (blue). Speed and regularity of the oscillatory signals were quantified as a mean and standard deviation (SD) of inter-peak intervals (IPIs), time lengths between contiguous peaks of the filtered waveform. (**C**) Evaluation of oscillation regularity. Pooling all IPIs within a retention period (e.g. D5, 4300 –5500 ms) generates a distribution of their occurrences. Higher regularity of oscillatory signals is indexed by a smaller variance or SD of the IPI distribution.

Although many studies have investigated changes in power (amplitude) of neural waveforms, an increasing number of studies focused on periodic and aperiodic aspects of neural oscillatory signals during WM task (Tian et al., 2017; Donoghue et al., 2020). Here I analyzed three different measures of oscillatory signals (**Fig. 3B**); amplitude, speed, and regularity. Using the Hilbert transformation, I measured changes in amplitude as an envelope of the filtered waveform in a frequency band of interest (dotted blue line in **Fig. 3B**). On the other hand, the speed and regularity of oscillatory signals were quantified by the inter-peak interval (IPI) analysis (Noguchi et al., 2019). I first identified all peaks on the filtered EEG waveform, measuring IPIs as time lengths between contiguous peaks. A mean length of IPIs pooled over a given period (e.g. 4300 – 5500 ms in case of delay 5) indexes a speed of neural oscillations, because slow/fast oscillatory signals produce longer/shorter IPIs. The regularity of neural waveforms, in contrast, was measured as a variance or standard deviation (SD) of IPIs. As shown in **Figure 3C**, irregular neural oscillations are characterized by a larger variance of IPIs, because such waveforms produce IPIs distant from the mean.

### Statistical procedures

An effect of semantic relatedness across MWs on neural activity was investigated by a comparison of Sem and NonSem trials. Of particular interest was the data in the delay 5 (D5) where a memory load became maximum. Avoiding visually-evoked potentials in response to 5th MW screen (4000 – 4200 ms), I compared the three measures of oscillatory signals at 4300 – 5500 ms between Sem and NonSem (**Fig. 4**). Since a paired *t*-test (N = 34 vs. 34) was repeated for 32 sensor positions, a problem of multiple comparisons was resolved by controlling false discovery rate (FDR). I adjusted a statistical threshold based on the Benjamini-Hochberg correction (Benjamini and Hochberg, 1995) with the *q*-value set at 0.05. Sensors showing a significant difference after this correction were marked with orange rectangles in **Figure 4** and **Figure 5**.

**Figure 4.**
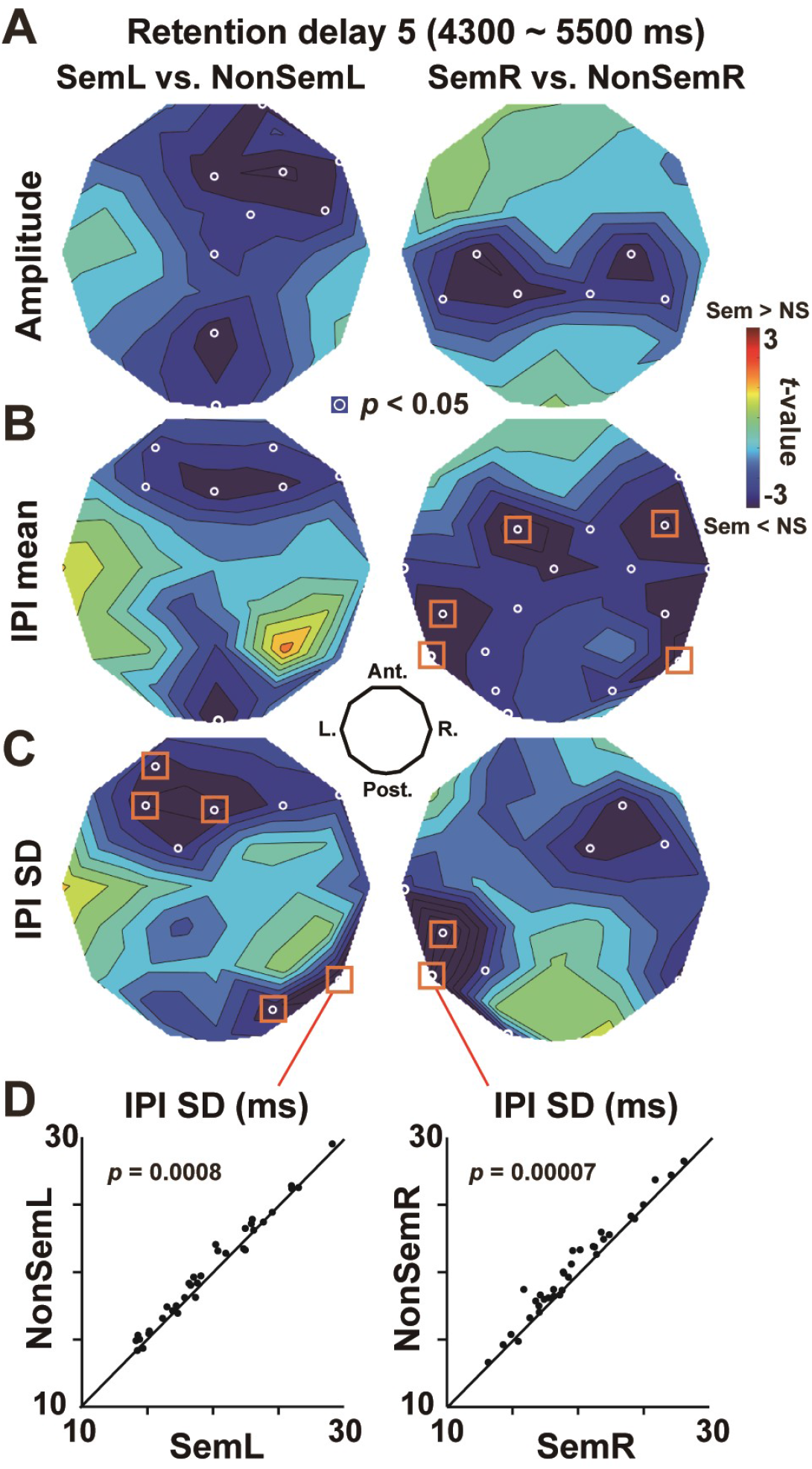
Effects of semantic relatedness on the three oscillatory measures. (**A**) *t*-map of oscillation amplitude (**Fig. 3B**, blue line). Mean amplitudes over 4300 – 5500 ms (delay 5) were compared between Sem and NonSem trials at each EEG sensor. Resultant *t*-values (negative: Sem < NonSem) were color-coded over the layout of 32 sensors. (**B**) *t*-map on mean IPIs (**C**) *t*-map on SDs of IPIs. White circles denote sensors showing a significant (*p* < 0.05, uncorrected) difference, while orange rectangles denote a significant difference after a correction of multiple comparisons. Memorizing 5 words semantically associated induced a reduction in SD of IPIs (EEG waveforms with higher regularity) over the temporal region contralateral to a cued hemifield (e.g. right temporal cortex in SemL). (**D**) Individual data of 34 participants at T6 and T5. The SD of IPIs in Sem (*x*) and NonSem (*y*) trials are shown on two-dimensional plots with a 45-deg line.

**Figure 5.**
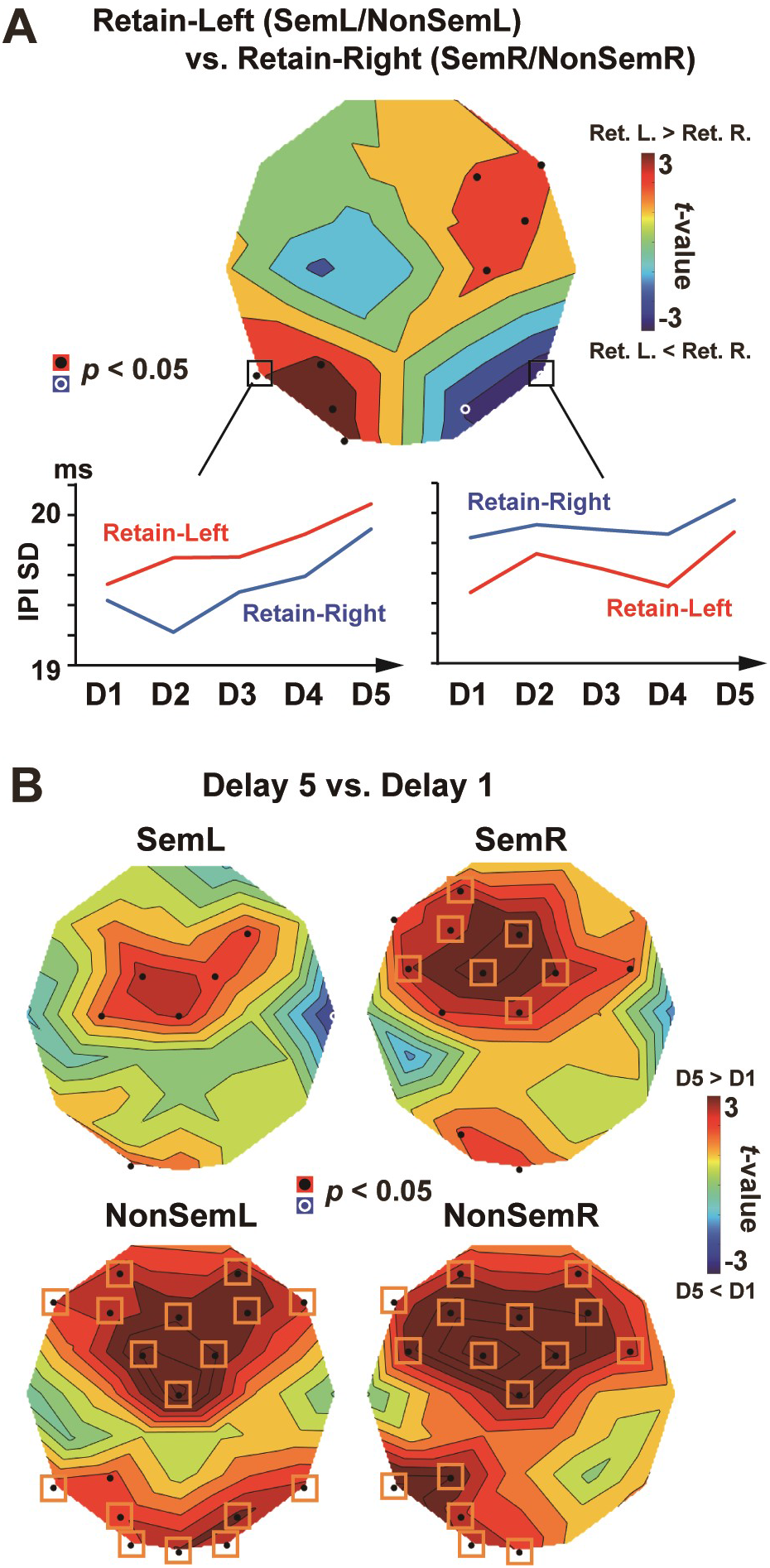
Changes oscillation regularity over 5 delays (D1 – D5). (**A**) *t*-map of IPI-SD (averaged across the 5 delays) between Retain-Left trials (SemL and NonSemL) and Retain-Right (SemR and NonSemR) trials. Attentive processing of words in a left/right hemifield induces a decrease in SD (increase in regularity) over the right/left hemisphere. One can see, however, a gradual increase in SD from D1 to D5, which presumably reflects an accumulating memory load over time. (**B**) *t*-map of IPI-SD between D5 (4300 – 5500 ms) and D1 (300 – 1000 ms). In NonSem trials (lower panels), the time-related increase in SD (D5 > D1, shown in red) was prominent in the frontal cortex and posterior regions contralateral to a cued hemifield. This increase in SD was inhibited by semantic relatedness across MWs (Sem trials, upper panels), especially in the posterior regions. Black dots denote sensors showing a significant (*p* < 0.05, uncorrected) difference, while orange rectangles denote a significant difference after a correction of multiple comparisons.

### Experiment 2

Results in Experiment 1 showed higher regularity of EEG waveforms in Sem than NonSem conditions (see **Results**). This suggested that semantically-related words induced similar patterns of neural oscillations that were easy to be integrated when co-stored in vWM (**Fig. 1A**). I examined this point more directly in Experiment 2. Specifically, the same set of 300 words as Experiment 1 were presented individually (one by one) in Experiment 2. A semantic-correlation matrix for each pair of words (300 × 300) were compared with another correlation matrix (300 × 300) for neural oscillatory responses to those words. If those two matrices are highly similar to each other, this would show an effect of semantic factor on a similarity of oscillatory responses, explaining why I observed the high-regularity signals in Sem trials of Experiment 1.

Each participant in Experiment 2 performed two tasks (**Fig. 6**). The first task involved a memory of five words sequentially presented (**Fig. 6A**). This was identical to the vWM task in Experiment 1, except that there was no attentional direction by the cue (a MW screen in Exp. 2 had only one word in its center). Participants underwent 2 sessions of 60 trials in which Sem and NonSem trials were intermixed in a random order. There were three types of probes in Sem condition (20 trials with old probes, 20 trials with new probes, and 20 trials for lure probes) but two types of probes in NonSem condition (30 trials with old probe and 30 trials with new probes). In the second task, the same set of 300 words as Experiment 1 was presented one by one (**Fig. 6B**). Participants performed an animacy judgment task on each word, pressing one key to animate and another to non-animate objects (150 trials × 2 sessions). An experiment started with the animacy judgment task, followed by the memory task. Other details (measurements of EEG data and analyses of behavioral data) were identical to Experiment 1.

**Figure 6.**
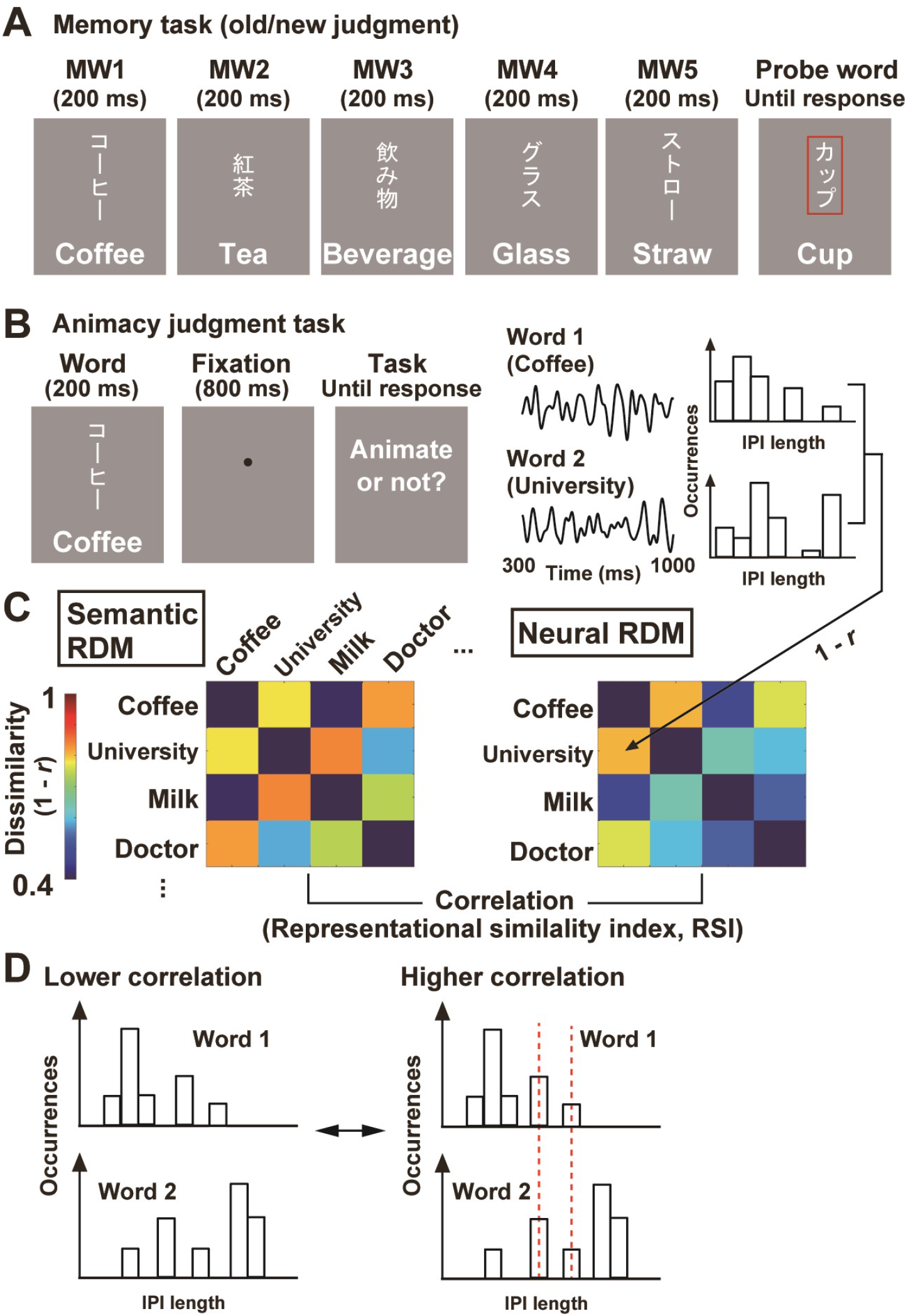
Experiment 2. (**A**) Memory task. Basic structures were the same as Experiment 1, except that each MW screen had only one word in its center. (**B**) Animacy judgment task. The same set of 300 words as the memory task was presented individually. Participants pressed one key to a word representing an animate object and pressed another to a non-animate object. (**C**) Representational similarity analysis (RSA). Using semantic vectors, I constructed a representational dissimilarity matrix (RDM, 300 × 300) showing a semantic distance (1 – *r*) for each pair of words (semantic RDM). I also made a neural RDM (300 × 300) based on a correlation of IPI histograms in response to each word in the animacy-judgment task. A high RSI (representational similarity index, a correlation between the semantic and neural RDMs) indicates that semantically-associated words induce similar distributions of IPIs at that EEG sensor. (**D**) The correlation of IPI histograms between two words. EEG waveforms in alpha-to-beta band (8 - 30 Hz) produced about 15 IPIs in a time window of analysis (300 – 1000 ms after a word onset). A higher correlation would be observed when the two histograms share the same set of neural oscillations (IPIs).

### Representational similarity analysis

EEG data in the animacy judgment task were used for the representational similarity analysis or RSA (Kriegeskorte and Kievit, 2013; Liu et al., 2020; Liu et al., 2021). First, I made a representational dissimilarity matrix (RDM) reflecting a semantic distance for each pair of 300 words (**Fig. 6C**). Each cell in this semantic RDM (dissimilarity index or DI) was defined as 1 - *r*, where *r* was a correlation between semantic vectors of two words. Next, I made another RDM based on a correlation of IPIs (neural RDM). A histogram of alpha-to-beta IPIs at 300 – 1000 ms was depicted for each word, with its vector defined as

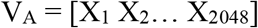

where V_A_ denotes an IPI vector of word A, and X_1_ – X_2048_ shows numbers of IPIs at bin 1 (0.488 ms) to bin 2048 (1000 ms). Each DI in the neural RDM was 1-*r*, where *r* was a correlation between IPI vectors of two words (e.g. V_A_ vs. V_B_). As shown in **Figure 6D**, a higher correlation would be observed when two histograms share the same set of neural oscillations (IPIs).

Finally, a correlation between semantic and neural RDMs was computed at each EEG sensor (representational similarity index or RSI). Since bottom-left entries were identical to top-right entries in each RDM, I compared the bottom-left halves of the semantic and neural RDMs, excluding diagonal components (0). A high RSI indicates that semantically-associated words produced similar distributions of IPIs. Statistical significance of those RSI was evaluated through a comparison with random data. By permutating the word labels of semantic RDM randomly for 1000 times, I generated a distribution of RSIs under a hypothesis of null effect. Rarity (*p*-value) of RSI in actual data was estimated as its percentile in this null distribution. A problem of multiple comparisons over the 32 sensors was resolved by the FDR-controlling approach as Experiment 1.

As a control, I tested whether distributions of alpha-to-beta IPIs were modulated by visual (not semantic) factors of word stimuli. A new RDM reflecting visual dissimilarities of word pairs was made for this analysis (visual RDM, **Fig. 7C**). The 300 words in the present study consisted of two types of Japanese letters: Kana (phonograms) and Kanji (ideograms, imported from China). The Kanji letters are characterized by higher spatial frequency and complexity than Kana letters (Horie et al., 2012). I thus classified the 300 words into three categories; Kana words, Kanji words, and mixtures of Kana and Kanji. The DI in the visual RDM was set at 1 when two words belonged to the same category and at 3 when one was a Kana word and the other was a Kanji word. A DI of 2 was given to a word pair of a Kana-Kanji mixture and a Kana/Kanji word. A rarity map of RSIs between the visual and neural RDMs was then computed and shown as the right panel in **Figure 7C**.

**Figure 7.**
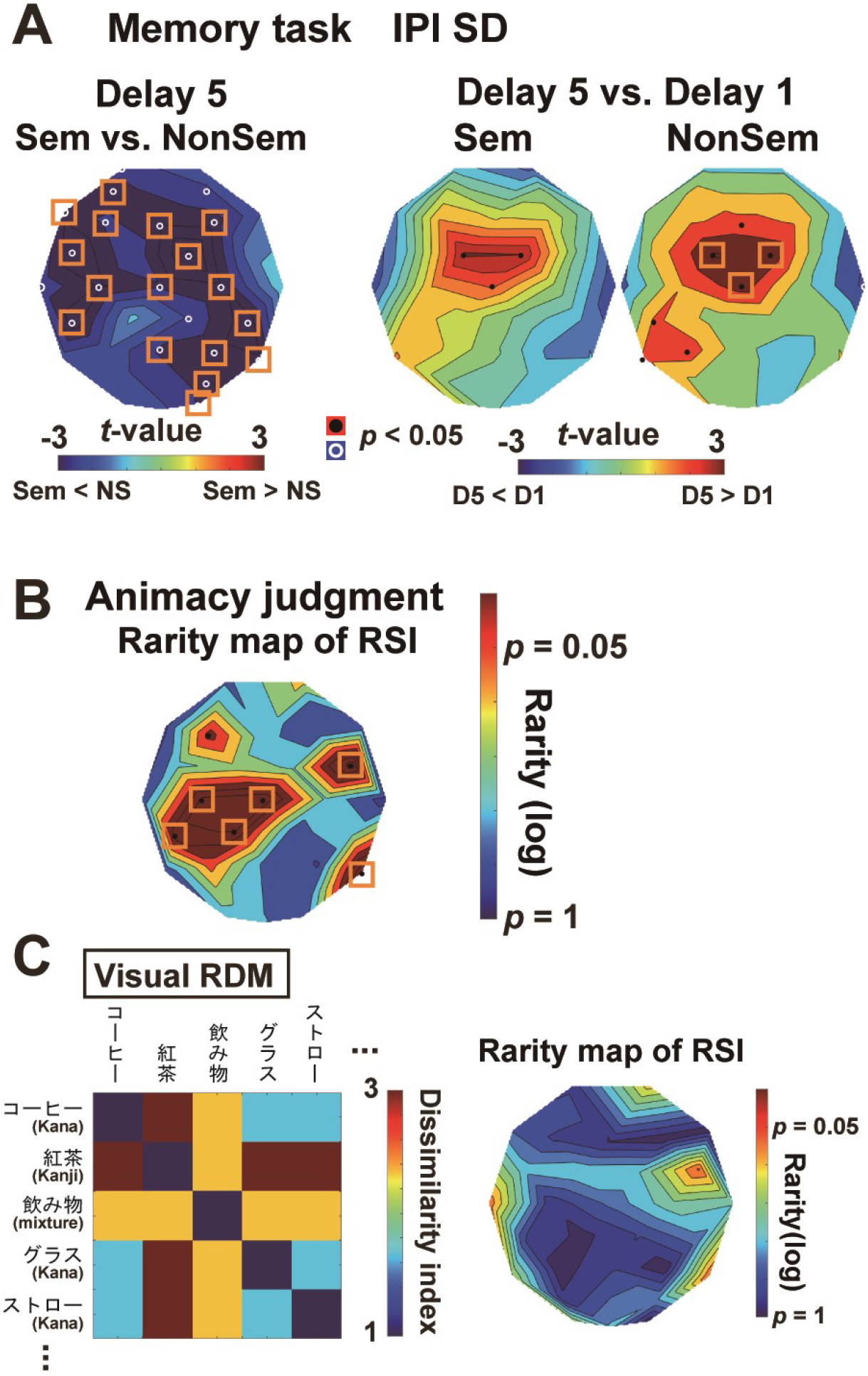
Results of Experiment 2. (**A**) Memory task. The SD of IPIs during D5 was found to be lower in Sem than NonSem trials (left panel). Semantic associations across five MWs in Sem trials mitigated time-related increase in SD (D1 < D5) over the left temporal regions (right panels). Those results replicated Experiment 1. (**B**) A rarity map of RSI in animacy judgment task. Rarity (*p*-value) of RSI in actual data was estimated through a random permutation of semantic RDM for 1000 times (see text for details). Orange rectangles show sensors with significant RSI after a correction of multiple comparisons. (**C**) Control. A new RDM based on visual similarity between two words (visual RDM, left panel) was compared with neural RDM in **Figure 6C**. No significant RSI was observed in the rarity map (left panel), indicating that correlations of IPI histogram were not modulated by visual factors.

## Results

### Behavioral data (Experiment 1)

**Figure 2C** shows the d-prime (*d’*) of the old/new judgment task. I observed the *d’* significantly higher in SemL (3.48 ± 0.08, mean ± SE across participants) than NonSemL (3.03 ± 0.15) trials (*t*(33) = 3.34, *p* = 0.002, *d* = 0.63), showing that semantic relatedness among MWs facilitated a retention of those words when they were presented in left visual field (right hemisphere). No significant difference, in contrast, was seen (*t*(33) = 1.05, *p* = 0.30, *d* = 0.20) when participants memorized words in right visual field (SemR: *d’* = 3.56 ± 0.07, NonSemR: *d’* = 3.45 ± 0.11).

**Figure 2D** displays FA rates to new (unseen) probes and lure probes. In SemL, the FA rates to lures (4.56 ± 0.93 %) were significantly higher (*t*(33) = 4.04, *p* = 0.0003, *d* = 0.70) than those to new words (1.18 ± 0.70 %), showing a false memory arising from a semantic integration. Similar results were observed in SemR (lures: 4.26 ± 0.97 %, new words: 0.29 ± 0.29 %, *t*(33) = 4.34, *p* = 0.0001, *d* = 0.95)

Taken together, those data indicated that participants integrated semantic information of MWs irrespective of whether they were presented in left or right visual field. A left-hemispheric dominance of vWM (Smith and Jonides, 1997; Nagel et al., 2013), however, might obscure a difference in *d’* between SemR and NonSemR.

### EEG data

**Figure 3A** shows a time-frequency power spectra over the left temporal cortex in SemR. Prominent power changes during the five retention periods (D1 - D5) were observed in alpha-to-beta band (8 – 30 Hz). These data were consistent with mounting evidence from previous studies showing an involvement of alpha (Wianda and Ross, 2019), beta (Weiss and Mueller, 2012; Sato et al., 2021), and alpha-to-beta (Meltzer et al., 2017; Mapelli and Ozkurt, 2019; Proskovec et al., 2019) rhythms in semantic processing and WM. I thus mainly focused on EEG signals in the alpha-to-beta band (8 – 30 Hz) below.

Comparisons of the three oscillatory measures (amplitude, speed, and regularity) during D5 between Sem and NonSem trials are shown in *t*-maps of **Figure 4**. Although amplitudes of EEG waveforms at 8-30 Hz tended to be smaller in Sem than NonSem trials, no significant difference was observed after a correction of multiple comparisons (**Fig. 4A**). The *t*-map of mean IPI, a measure for oscillation speed, showed significant differences (Sem < NonSem) over frontal and temporal regions (**Fig. 4B**). Although these data suggested an acceleration of brain rhythm caused by a semantic integration, the change in mean IPI was limited to when participants memorized items in right visual field (SemR vs. NonSemR, right panel). Finally, the SD of IPIs, a measure for irregularity of EEG waveforms, showed significant reductions in Sem compared to NonSem trials (**Fig. 4C**) both when memory items were presented in left and right visual fields. Clear differences were found over the posterior temporal cortex contralateral to a cued hemifield (**Fig. 4D**). The same results were obtained when I compared the Fano factor, a variance of IPIs normalized to their mean, between Sem and NonSem trials (**Fig. S1 in Supplementary materials**). Those results indicate an increase in regularity (decrease in IPI-SD) of oscillatory signals induced by semantic relatedness of MWs. Indeed, the reduction in IPI-SD between Sem and NonSem were significantly correlated with the FA rates to lure probes (**Fig. S2**), suggesting a close relationship of IPI-SD with a false memory (semantic integration).

### Time-related changes in regularity of oscillatory signals

More detailed information about SDs of IPIs are provided in **Figure 5**. First, a comparison between Retain-Left trials (SemL and NonSemL) and Retain-Right trials (SemR and NonSemR) showed positive and negative *t*-values in left and right occipito-temporal cortex, respectively (**Fig. 5A**). Attentive processing of words in left/right hemifield therefore induced a reduction of IPI-SD (increase in regularity) over the right/left hemisphere.

This neural signature of information processing (high regularity), however, diminished over time. One can see a gradual increase in IPI-SD from D1 (300 – 1000 ms) to D5 (4300 – 5500 ms), presumably reflecting an accumulating memory load. **Figure 5B** shows *t*-maps of IPI-SD between D1 and D5. In NonSem trials (lower panels), time-related increase in SD (D5 > D1, shown in red) was prominent in the posterior regions contralateral to a memory field. This increase in SD was mitigated in Sem trials (upper panels), indicating that semantic integration of MWs inhibited a generation of irregular components in the posterior regions.

### Results in Experiment 2

Means and SEs of *d’* in the memory task were 3.56 ± 0.09 in Sem and 3.74 ± 0.12 in NonSem trials. No significant difference was observed (*t*(26) = 2.01, *p* = 0.055, *d* = 0.32). The FA rate to lure probes (5.56 ± 1.72 %) was significantly higher (*t*(26) = 2.51, *p* = 0.019, *d* = 0.55) than that to new probes (1.85 ± 0.66 %). These behavioral results were similar to those in Retain-Right trials in Experiment 1. The *t*-maps of IPI-SD were shown in **Figure 7A**. Oscillatory signals during D5 were more regular in Sem than NonSem trials (left panel). The time-related increase in IPI-SD was inhibited in Sem trials over the left temporal cortex. Those data replicated Experiment 1.

Mean and SE of accuracy in the animacy judgment task was 99.28 ± 0.28 %. A rarity map of RSI between the semantic and neural RDMs is shown in **Figure 7B**. Significant RSIs were found in the frontal, parietal, and temporal regions especially in the left hemisphere. Results of RSA separately conducted for 34 words representing animate object and 266 words representing non-animate objects are provided in **Figure S3**. The RSI over the left temporal regions were significant in all analyses. These data showed that semantically-associated words induced similar sets of IPIs, which were consistent with the high regularity of oscillatory signals when co-stored in vWM (**Fig. 7A**).

I performed another RSA using a RDM reflecting visual (not semantic) similarities of the 300 words (**Fig. 7C**). Non-significant RSI in this analysis (right panel) indicated that visual factors had no effect on a distribution of alpha-to-beta IPIs at 300 – 1000 ms.

## Discussion

In the present study, I compared neural oscillatory signals when human participants retained the information of five words semantically related (Sem trial) or not (NonSem trial). Results revealed a reduced SD (increased regularity) of IPIs in Sem than NonSem conditions over the temporal cortex contralateral to memory items (Exp.1). The reduction of mean IPIs (an acceleration of brain rhythm) was also observed when memory items were presented in right visual field (left hemisphere). In Experiment 2, I presented the same set of words individually in a non-memory (animacy judgment) task, finding that semantically-related words induced similar distributions of IPIs. These results can be explained by assuming that each semantic feature of a word is retrieved and retained as a neural oscillation at specific frequencies. Memorizing words with a common semantic feature would cause a resonance (or sharing) of the oscillatory signals across the words, resulting in increased regularity of EEG waveforms during the retention period (**Fig. 1**).

In both Experiments 1 and 2, key regions for a semantic integration were found over the posterior temporal lobe. This is consistent with recent evidence. For example, Volfart et al. (2021) reported neural activity in the posterior temporal cortex related to a semantic categorization (e.g. discriminating animal from city names) of visually-presented words (Volfart et al., 2021). In light of a previous literature, a source region for present data might be posterior middle temporal gyrus (pMTG) (Fairhall and Caramazza, 2013; Chen et al., 2017). In order to integrate the information of five words, the brain had to access semantic knowledge of each word and connect the words under a common concept. The pMTG is thought to play a critical role in such a process involving an immediate recall and re-structuring of semantic networks (Lambon Ralph et al., 2017).

Several previous studies have investigated an effect of across-word (or across-object) relationships on oscillatory signals in the human brain (Mapelli and Ozkurt, 2019). They, however, mainly analyzed changes in oscillatory powers, such as an alpha-power decrease during the retention of semantically-related words (Melnik et al., 2017) and a beta-power decrease induced by the sentence superiority effect (Bonhage et al., 2017). Consistent with those data, I found decreases in power of alpha-to-beta rhythm in Sem compared to NonSem trials (**Fig. 4A**). Replicating those previous findings, I further showed that semantic relatedness of words also modulated temporal measures (speed and regularity) of brain rhythms.

Previous studies have proposed cross-frequency couplings as a neural model of multi-unit memory. Typical example is the phase-amplitude coupling between theta and gamma rhythms (Heusser et al., 2016; Bahramisharif et al., 2018; Griffiths et al., 2021). In this model, individual memory items are represented by oscillatory signals in a gamma range. Information of multiple items is retained by sequentially activating their representations (gamma activities) on a theta cycle. Although this model based on the serial reactivation is potent as a neural substrate of multi-unit memory and has been supported by compelling evidence(Canolty et al., 2006; Sauseng et al., 2009; Long and Kahana, 2017; Kerren et al., 2018; Reinhart and Nguyen, 2019; Huang et al., 2021; Reddy et al., 2021), it does not mention explicitly how the information of items is integrated in the buffer of WM. Indeed, this line of studies have focused on how the brain segregates (not integrates) information of semantically-related concepts e.g. by reactivating them in distant phases of theta cycle (Kunz et al., 2019; Pacheco Estefan et al., 2021). The present results on a semantic integration would make up for the point those previous models have not fully addressed.

In conclusion, the present data provided an insight on how verbal information was represented and integrated as neural (electric) signals in the healthy human brain. Semantic chunking is known to be a key method to enhance one’s memory capacity. The current data might be also useful to develop a new method to prevent age-related degradation of visual and verbal WM (Lorenc et al., 2021).

## Acknowledgments

This work was supported by KAKENHI Grants Number 19H04430 from the Japan Society for the Promotion of Science (JSPS) to Y.N. I thank Nahomi Sato and Taeko Kaneda for their technical support. All data supporting the findings of this study are available from Y.N. upon reasonable request.

