## Supplemental Figures S1 - S3 for "Harmonic memory signals in the human cerebral cortex induced by sematic relatedness of words"

### Supplementary Materials (Fig. S1 - S3)

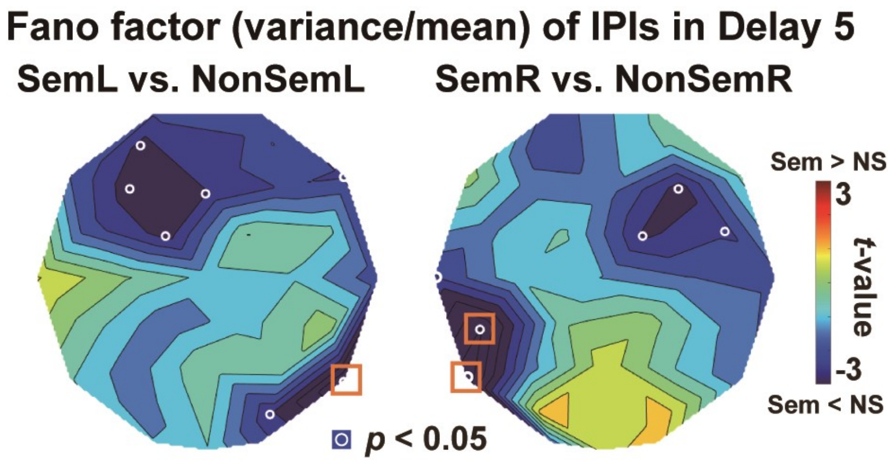


**Figure S1.** *t*-maps of Fano factor of IPIs. Fano factor is a variance of IPIs divided (normalized) by their mean and thus provides an unbiased measure for the irregularity of neural oscillation. Semantic relatedness of memory words induced reduction in Fano factor (increases in regularity) over the occipito-temporal cortex contralateral to a cued hemifield. Orange rectangles denote a significant difference after a correction of multiple comparisons.

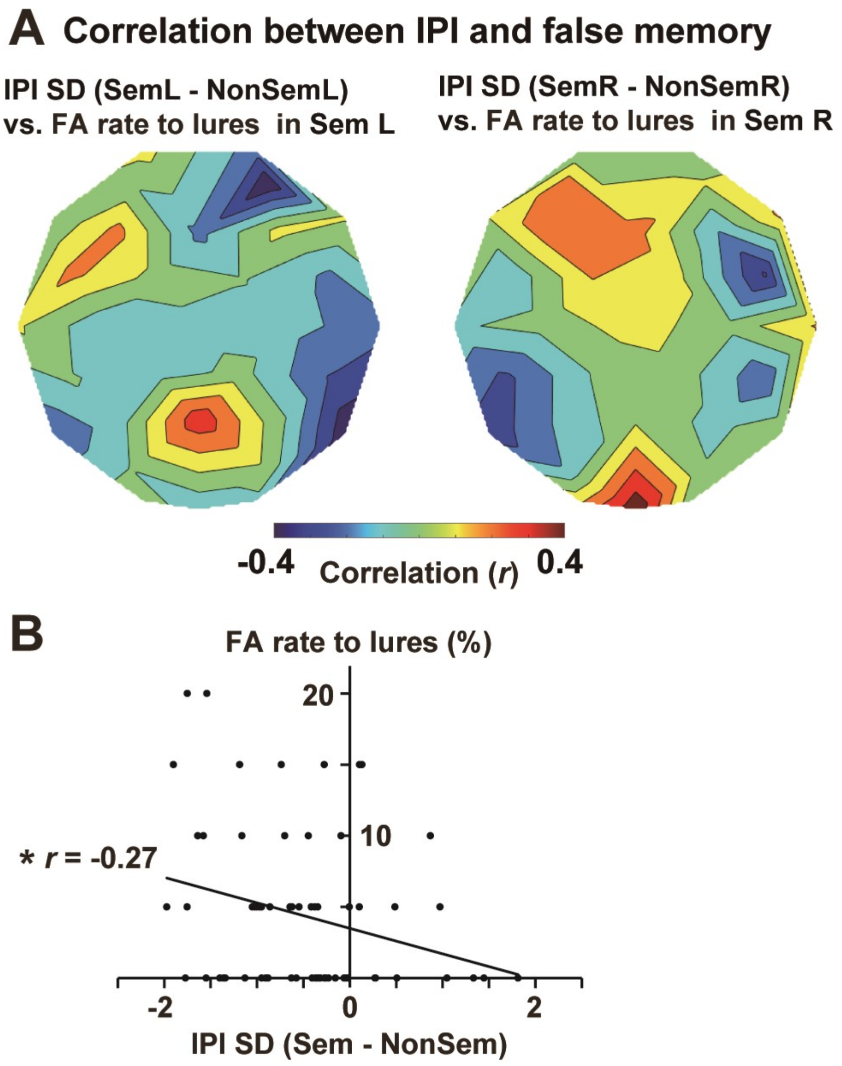


**Figure S2.** Correlation between IPI and behavioral data (false memory) in Experiment 1. (**A**) Correlation *r*-maps between a difference in IPI-SD (Sem - NonSem) and a false-alarm (FA) rate to lure probes. Negative correlations over the temporal cortex indicate that participants with a greater reduction of IPI-SD (Sem < NonSem) showed a higher rate of false memory. (**B**) Individual data. Based on results of **Figure 4C**, we selected four EEG sensors showing a significant reduction of IPI-SD in Sem compared to NonSem trials; PO4, T6, T5, and CP5. The IPI data in the right temporal cortex (averages of PO4 and T6) are plotted against FA rates in SemL (N = 34), and the IPI data in the left temporal cortex (averages of T5 and CP5) are plotted against FA rates in SemR (N = 34). *$p < 0.05$.

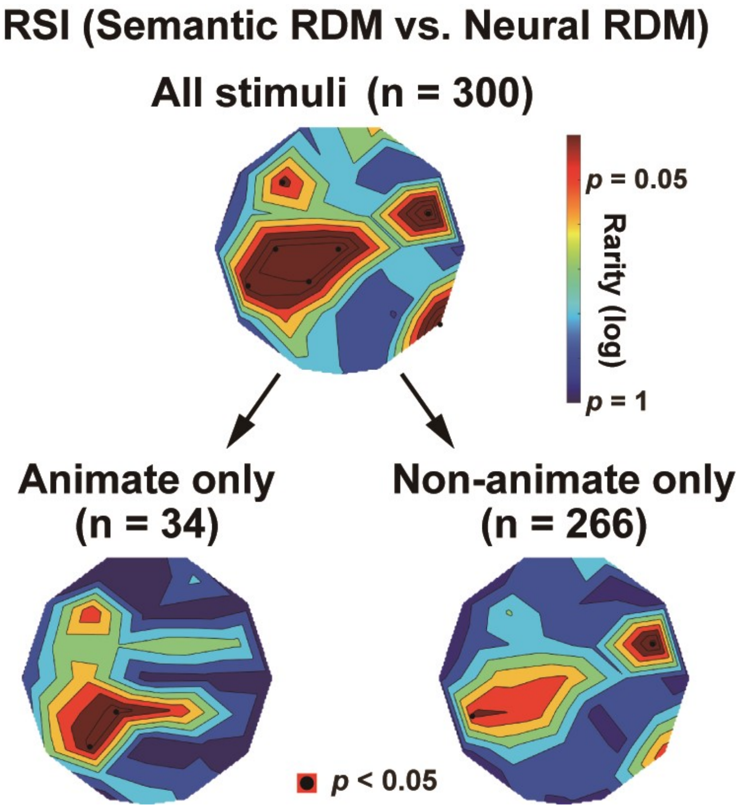


**Figure S3.** Representational similarity analysis (RSA) using a subset of words. Top panel: Rarity maps of RSI (representational similarity index) using the data of all 300 words (same as **Fig. 7B**). Lower-left panel: Results of RSA using the data of 34 words representing animate objects. A smaller size of semantic and neural RDMs (34 × 34) was used in this analysis. Lower-right panel: Results of RSA using the data of 266 words representing non-animate objects. Significant RSIs were observed over the left temporal regions in all maps. The RSI in **Figure 7B** thus reflect the similarity of IPIs modulated by a semantic distance for each pair of words, not emerging from a task requirement (a binary distinction between animate and non-animate objects).
